# Loss of *Prdm12* during development, but not in mature nociceptors, causes defects in pain sensation

**DOI:** 10.1101/2020.09.07.286286

**Authors:** Mark A. Landy, Megan Goyal, Katherine M. Casey, Chen Liu, Helen C. Lai

## Abstract

*Prdm12* is as a key transcription factor in nociceptor neurogenesis. Mutations of *Prdm12* cause Congenital Insensitivity to Pain (CIP) due to failure of nociceptor development. However, precisely how deletion of *Prdm12* during development or adulthood affects nociception is unknown. Here, we employ tissue- and temporal-specific knockout mouse models to test the function of *Prdm12* during development and in adulthood. We find that constitutive loss of *Prdm12* causes deficiencies in proliferation during sensory neurogenesis. We also demonstrate that conditional knockout from dorsal root ganglia (DRGs) during embryogenesis causes defects in nociception. In contrast, we find that in adult DRGs, *Prdm12* is dispensable for pain sensation and injury-induced hypersensitivity. Using transcriptomic analysis, we found unique changes in adult *Prdm12* knockout DRGs compared to embryonic knockout, and that PRDM12 is likely a transcriptional activator in the adult. Overall, we find that the function of PRDM12 changes over developmental time.

## Introduction

Nociception is a critical warning system for the detection of tissue damage by noxious stimuli, but it can often go awry, resulting in either greatly increased or decreased pain sensation. Patients with Congenital Insensitivity to Pain (CIP) are a prime example of the latter. These people cannot feel mechanical, thermal, or inflammatory pain, or even discomfort associated with internal injuries. Cases of CIP have typically been associated with mutations of nerve growth factor (NGF) or its receptor *Ntrk1* (TRKA), which cause failure of nociceptor development (Capsoni et al., 2011; Carvalho et al., 2011; Einarsdottir et al., 2004), or in the voltage-gated sodium channels, *Scn9a* (Nav1.7) or *Scn11a* (Nav1.9) (Cox et al., 2006; Leipold et al., 2013). More recently, additional mutations have been identified to cause CIP (Nahorski et al., 2015), including those of the transcription factor PRDM12 (PRDI-BF1-RIZ homologous domain-containing family) (Chen et al., 2015). As in other forms of CIP, patients with *Prdm12*-associated CIP are unable to feel pain due to noxious chemical, thermal, or mechanical stimuli, but retain normal touch, proprioception, and tickle sensations (Chen et al., 2015; Saini et al., 2017; Zhang et al., 2016). Therefore, *Prdm12* or its downstream effectors may serve as potential novel analgesic targets similar to drugs that have been developed targeting other genes underlying CIP, NGF and Nav1.7 (Hoffman et al., 2011).

The PRDM family of transcription factors are known to have essential roles in cell fate transitions (Hohenauer and Moore, 2012) and many lines of evidence suggest that PRDM12 has an essential role in nociceptor development and maintenance. It is highly conserved from mouse to human, with 98% protein identity, suggesting a highly conserved function, and opening the gateway for study in mouse models. Indeed, consistent with the idea of having a highly conserved function, loss-of-function of PRDM12 in humans or its homologs in *Drosophila*, frog embryos, and mice leads to abnormalities in sensory neuron development (Bartesaghi et al., 2019; Chen et al., 2015; Desiderio et al., 2019; Moore et al., 2004; Nagy et al., 2015). Additionally, *Prdm12* is specifically expressed in myelinated Aδ- and unmyelinated C-fiber nociceptors into adulthood (Chen et al., 2015; Kinameri et al., 2008; Matsukawa et al., 2015; Nagy et al., 2015; Sharma et al., 2020; Thelie et al., 2015; Usoskin et al., 2014). The highly conserved sequence, function, and expression pattern raise the possibility that PRDM12 is serving an important role in nociceptor biogenesis.

Structurally, PRDM12 consists of a PR domain, three zinc finger domains, and a polyalanine repeat. The PR domain is characteristic of all members of the PRDM family of proteins, and bears weak homology to SET domains, which have histone methyltransferase (HMT) activity (Kinameri et al., 2008). However, PRDM12 itself is reported to have weak endogenous HMT activity, and is thought to exert repressive activity predominantly through interaction with EHMT2 (euchromatic histone-lysine N-methyltransferase, also called G9a), which catalyzes repressive chromatin marks (Yang and Shinkai, 2013). This interaction was shown to be dependent on the second zinc finger domain (ZnF2), contained in exon V (Yang and Shinkai, 2013).

Thus far, most of the work surrounding *Prdm12* has focused on its role in nociceptor neurogenesis. Early reports indicated that PRDM12 promoted expression of sensory neuronal markers (Kinameri et al., 2008; Matsukawa et al., 2015; Thelie et al., 2015; Yang and Shinkai, 2013). Consistent with this, *Prdm12* was found to be necessary for the initiation and maintenance of tropomyosin receptor kinase A (TRKA) expression, a marker for early nociceptor development (Desiderio et al., 2019). In addition, in the absence of *Prdm12*, the entire nociceptor lineage failed to develop. However, the mechanism by which *Prdm12* knockout leads to a deficiency in nociceptors is unclear. Work by Bartesaghi et al. found evidence for decreased proliferation in *Prdm12* knockout mice with no change in cell death (Bartesaghi et al., 2019), while work from Desiderio et al. found the opposite, that there was no change in proliferation, but there was an increase in cell death (Desiderio et al., 2019). Differences in the way proliferation and cell death were quantitated in these studies could account for these discrepancies.

Therefore, in our study, we sought to further clarify the mechanism by which *Prdm12* controls nociceptor development. Furthermore, we wanted to examine the behavioral defects in mice that lack *Prdm12* during embryogenesis and determine whether it is an important component of pain sensation in mature sensory neurons. To do this, we generated three mouse models with which to study the effect of *Prdm12*-knockout at different timepoints: (1) *Prdm12*^*-/-*^, a constitutive knockout to assess the early embryonic changes resulting from *Prdm12* deletion, (2) *Prdm12*^*AvilCKO*^, a dorsal root ganglion (DRG)-specific conditional knockout to assess pain sensation in mice lacking *Prdm12* from around E12.5 onwards, and (3) *Prdm12*^*AvilERT2CKO*^, a tamoxifen-inducible DRG-specific conditional knockout to investigate the role of *Prdm12* in adult nociceptors. With these models, we confirm that *Prdm12* expression is necessary for the development of nociceptors, and show that its absence results in defects in proliferation during neurogenesis. Furthermore, we demonstrate that embryonic sensory neuron-specific knockout of the gene results in mice with defects in mechanical and cold nociception, as well as itch. Finally, we show that knockout of *Prdm12* in mature DRGs does not impact nociception, even under conditions of neuropathic injury or inflammation. However, we provide transcriptomic evidence for an alternate function of *Prdm12* in these neurons compared to embryonic development and that it is potentially an activating transcription factor in the adult rather than a repressor.

## Materials & Methods

### Mouse strains

The following mouse strains were used: *Prdm12*^*F/F*^ (Chen et al., 2020), CAG-Cre (Sakai and Miyazaki, 1997), *Advillin*^*Cre/+*^ (JAX#032536) (Hasegawa et al., 2007), *R26*^*LSL-tdTomato/+*^ (Ai14, JAX#007908) (Madisen et al., 2010), *Avil*^*CreERT2*^*BAC* (JAX#032027) (Lau et al., 2011), *R26*^*LSL-EYFP/+*^ (Ai3, JAX#007903). All mice were outbred and thus are mixed strains (at least C57Bl/6J, C57Bl/6N, and ICR). Both male and female mice were used for all studies. No sex differences were noted for any quantified data, therefore sexes of the same genotype were pooled for analysis. Mice crossed to *Advillin*^*Cre/+*^ always included a fluorescent reporter and were screened for “dysregulated” expression. Normal fluorescence is visible in the trigeminal and dorsal root ganglia of pups within the first three days after birth, while dysregulation results in patchy fluorescence of the whole body. Mice expressing *Avil*^*CreERT2*^*BAC* were injected with tamoxifen (Sigma) at eight weeks of age. Injections were given over a period of five days (1/day, 120mg/kg delivered intraperitoneally (i.p.) as a 40mg/ml solution dissolved in sunflower oil with 10% ethanol) (Lau et al., 2011; Sikandar et al., 2018). All animal experiments were approved by the Institutional Animal Care and Use Committee at UT Southwestern.

### Behavior assessments

For all behavioral tests, animals were habituated for 30 min one day before testing, and again immediately prior to testing. A single experimenter conducted all tests and was blinded to genotype. The subsequent statistical analyses included all data points; no methods were used to predetermine sample sizes. Littermates were used as controls.

### von Frey mechanical Sensitivity

von Frey withdrawal thresholds were determined using the simplified up-down method (Bonin et al., 2014). After acclimation in plastic chambers with wire mesh flooring, mice were tested with graded filaments from 0.008 to 2.0 g applied for ∼3 sec to the plantar hindpaw with at least 5 min between each application. Responses on both the left and right paw were recorded. Toe spreading, flinching, or licking was recorded as a positive response.

### Tail clip and Pinprick

Mechanical nociception was assessed using the tail clip and pinprick assays. For tail clip, electrical tape was wrapped around the jaws of a 1 cm binder clip, which was then attached to the tail, about 1 cm from the rostral end. The latency to response (biting or clawing at the clip, or otherwise trying to remove it) was recorded for a single trial, with a cutoff of 10 seconds. Pinprick was performed using 0.2 mm insect pins (FST 26002-20) to deliver a sharp mechanical stimulus. Mice were again acclimated in plastic chambers, then challenged 10 times on each paw, with 10 min between each trial. Positive responses (paw flinching, licking, or vocalization) was recorded and reported as a percentage of total trials.

### Rodent Pincher

The inflamed paw in the CFA inflammation model was tested with the Rodent Pincher Analgesia Meter (Bioseb). Mice were restrained by wrapping the mouse inside a paper towel with the inflamed paw exposed. The pincher was used to apply ramping pressure until a response (paw withdrawal or flicking, or vocalization) was observed. Three recordings were made per mouse, spaced at least 10 min apart.

### Heat sensitivity (hot plate and Hargreaves)

For hot plate, mice were placed directly on the plate (IITC) set to the designated temperature. The latency to response (hindpaw licking or flicking, or jumping) was recorded and averaged over three trials. Cutoff times were used to prevent injury as follows: 1 min for 50°C, 45 sec for 52°C, and 30 sec for 55°C. For Hargreaves, mice were acclimated on a heated (30°C) glass surface (IITC), then exposed to a beam of radiant heat following the standard Hargreaves method. Beam intensity was adjusted to result in latency of ∼10 sec in wildtype animals. Paw withdrawal latency was recorded for 3 exposures per paw, with at least a 5 min interval between exposures. A cutoff time of 30 seconds was used to prevent tissue damage.

### Cold sensitivity assays

Cold nociception was measured using either the cold plate or cold plantar assay. For cold plate, a cooling block was chilled at −20°C, then allowed to warm until the surface temperature reached 0°C as measured by an infrared thermometer. Mice were placed on the plate, and the latency to response (hindpaw licking or flicking, or jumping) was recorded and averaged over three trials. A cutoff time of 60 seconds was used to prevent injury. The cold plantar assay was performed using dry ice loaded into a syringe to stimulate the hindpaw (Brenner et al., 2012). Mice were placed on a thin, single pane of glass, and the tip of the dry ice pellet was pressed against the glass under the hindpaw. Withdrawal latency was recorded for 3 exposures per paw, with at least 5 min interval between exposures. A cutoff time of 30 seconds was used to prevent tissue damage.

### Itch assays

Itch sensation was measured by pruritogen injection into the nape of the neck (Kuraishi et al., 1995; Shimada and LaMotte, 2008). The injection area was shaved one day prior to testing. On the day of testing, mice were habituated in cylindrical containers for 30 minutes, then injected with 20 μL of histamine (100 μg/μL) or chloroquine (200 μg/μL) dissolved in PBS. The mice were video recorded for 30 min following pruritogen injection, and the videos were subsequently scored to determine total time spent scratching the injected area.

### Capsaicin test

The capsaicin test was performed by intraplantar injection to one hindpaw of 10 μL containing 0.3 μg/μL capsaicin (Sigma M2028) in 0.9% saline/10% ethanol/10% Tween-20 following acclimation. Mice were then video recorded for 10 minutes, and the videos were subsequently scored to determine time spent licking the injected paw.

### Touch assays

Non-nociceptive touch sensation was measured using the dynamic brush and sticky tape assays. For dynamic brush assay, were again acclimated in von Frey chambers. The tip of a cotton tipped applicator (Henry Schein) was teased apart to “fluff” it up and ensure no filaments were sticking straight up. The swab was then lightly brushed across the plantar surface of the hindpaw (about 1 s from heel to toe) 10 times per paw, with 10 min between each trial. Positive responses (paw flicking or withdrawal) are reported as a percentage of total trials. For sticky tape, a 5 mm x 5 mm piece of lab tape (Fisher) was adhered to the plantar surface of the hindpaw, and the mouse was allowed to freely explore its enclosure. Latency to removal of the tape was recorded and averaged across two trials per paw.

### Injury and inflammation

SNI surgery was performed as previously described (Decosterd and Woolf, 2000). Briefly, under 3% isoflurane anesthesia, the left sciatic nerve was exposed where it branches into the sural, tibial, and common peroneal nerves. The tibial and common peroneal nerves were tightly ligated using 5.0 silk suture, and a ∼3 mm section of nerve was removed just distal to the knot. Mice were allowed to recover for at least 48 hours prior to testing. For CFA-induced inflammation, 20 μL of Complete Freund’s Adjuvant (Sigma F5881) was injected into the plantar surface of the left hindpaw. Mice were first tested 6 hours post-injection to allow for development of the inflammatory response.

### Tissue processing

Pregnant dams were injected with 0.5 mg/mL EdU (5-ethynyl-2’-deoxyuridine, Carbosynth) at a dose of 10 μg EdU/g mouse 30 minutes prior to CO2 euthanasia for collection of embryos (Wang et al., 2011). Embryos fixed in 4% paraformaldehyde (PFA) in PBS for 2 hours at 4°C, washed overnight in PBS, and cryoprotected in 30% sucrose. Adult mice were anesthetized with Avertin (2,2,2-Tribromoethanol) (0.025 – 0.030 mL of 0.04 M Avertin in 2-methyl-2-butanol and distilled water/g mouse) and transcardially perfused, first with 0.012% w/v Heparin/PBS and then 4% PFA/PBS. A dorsal or ventral laminectomy exposed the spinal cord and DRGs for a post-fix in 4% PFA (2 hours at 4°C). Tissue was then washed in PBS and cryoprotected in 30% sucrose before the laminectomy was performed on the reverse side, allowing DRGs to be removed and embedded in OCT (Tissue-Tek Optimal Cutting Temperature Compound). All tissue was sectioned using a Leica CM1950 cryostat.

### Immunohistochemistry (IHC) and confocal imaging

Cryosections (20-30 μm were blocked with PBS/1% normal goat or donkey serum (Jackson labs)/0.3% Triton X-100 (Sigma) for up to 1 hr at room temperature (RT), then incubated overnight with primary antibody at 4°C. Sections were washed 3 times in PBS, then the appropriate secondary antibody (Alexa 488, 567, and/or 647, Invitrogen) was incubated for an hour at RT, and sections were again washed 3 times in PBS. For development of EdU signal, sections were then re-permeabilized in 0.5% Triton X-100 for 30 min at RT, then incubated in EdU detection solution (100 mM Tris pH 7.5, 4 mM CuSO4, 100 mM sodium ascorbate, 5 μM sulfo-Cy3 azide (Lumiprobe)) for 30 min at RT, and rinsed 3 times in PBS. Slides were mounted with Aqua-Poly/Mount (Polysciences Inc.), and coverslipped (Fisher). The following primary antibodies and dilutions were used: mouse anti-Islet1/2 (1:20,000; DSHB 39.4D5), goat anti-TRKA (1:20; R&D Systems AF1056), rabbit anti-RUNX3 (1:50,000; gift from Thomas Jessell), rabbit anti-CASP3 (1:50; BD Pharmingen 557035), IB4-488 (1:500, Invitrogen I21411), rabbit anti-CGRP (1:1000; Immunostar 24112), rabbit anti-NF200 (1:500; Sigma N4142), rabbit anti-TRPV1 (1:500; Alomone ACC-030).

Fluorescent images were taken on a Zeiss LSM880 confocal microscope with a 3 μm optical slice and the 20x objective. Images were pseudocolored with a magenta/yellow/cyan color scheme using Adobe Photoshop (Adobe) or Fiji. Cell counts were conducted manually using the built-in cell counter plugin on Fiji.

### In situ hybridization

A probe for ISH was generated targeting a 482 bp sequence entirely within exon V of *Prdm12*. ISH was performed as per standard protocols. Detailed protocol is available upon request. Briefly, DRG sections (30 μm) were dried at 50°C for 15 min then fixed in 4% PFA in DEPC-PBS for 20 min at RT. The sections were washed in DEPC-PBS for 5 min at RT then incubation in RIPA buffer (150 mM NaCl, 1% NP-40, 0.5% Na deoxycholate, 0.1% SDS, 1 mM EDTA, 50 mM Tris pH 8.0) for 60 min. The sections were then washed in DEPC-water followed by acetylation (500 µL of acetic anhydride in 200 mL of 0.1 M RNase-free triethanolamine-HCl at pH 8.0), washed in DEPC-PBS for 5 min., and prehybridized for 4 h at 64°C. Sections were incubated overnight at 64°C with 1–2 ng/µL of fresh *Prdm12* probe. The following day, a series of low and high stringency washes in 2x and 0.2X SSC as well as treatment with RNaseA and RNase T1 were performed. The sections were blocked in 10% inactivated sheep serum for 1 h followed by overnight incubation with 1:1000 anti-digoxygenin (DIG) antibody (Roche). The sections were washed in PBT and incubated with NBT/BCIP (Roche) staining solution until the blue precipitate formed. The slides were then washed in PBS and coverslipped with Aqua-Poly/Mount (Polysciences Inc.) mounting media. If ISH was followed by IHC, the sections were placed in PBS and then immunostained following the IHC protocol described above.

The RNAscope Fluorescent Multiplex Assay (Advanced Cell Diagnostics Inc., Hayward, CA) was performed according to the manufacturer’s instructions using a *Prdm12* exon V-specific probe (ACDBio, 320269-C3). All incubation steps were performed in a HybEZ™ II oven set to 40°C. The slides were washed with distilled water three times and incubated with Protease III for 40 sec. Slides were then washed with distilled water three times and incubated with the probe targeting *Prdm12* for 2 hours. The slides were washed two times thoroughly using 1X wash buffer for 2 min, then incubated with Amp 1-Fl for 30 minutes. The same process (washing then treatment) was repeated for Amp 2-Fl, Amp 3-Fl and Amp 4-Fl for 15, 30 and 15 minutes, respectively. Slides were washed three times in PBS for 10 minutes and coverslipped with Aqua-Poly/Mount (Polysciences, Inc.) mounting media.

### Microarrays

RNA was extracted from lumbar DRGs 2-5 following the manufacturer’s protocol with a Direct-zol RNA Miniprep Plus kit (Zymo R2071). RNA libraries were prepared and sequenced by the UTSW Microarray core facility on an Illumina NextSeq SE-75 sequencer at 40 million reads/sample. RNA-seq reads were mapped to mouse genome (mm10) and junctions were identified using tophat(v2.1.2) (Kim et al., 2013). Differential expression analysis was performed using cufflinks (v.2.2.1) (Trapnell et al., 2013). Both alignment and differential expression analysis were performed using default parameters. From differential expression results, genes showing expression of >=1 FPKM in either of the conditions cutoff was used in addition to FDR and fold change cutoffs.

### Experimental design and statistical tests

Cell counts were averaged across sections from 3 unique lumbar (L2-L5) DRGs per specimen, with 2-3 embryos or mice per timepoint or condition as indicated. Where applicable, counts were taken as a fraction of DRG area (measured in Fiji), or as a percentage of TOM^+^ cells. For both cell counts and behavior assessment, statistical analysis was conducted with the student’s t-test for pairwise comparisons, or with a 2-way ANOVA when analyzing data over time. All data and graphs were processed in Microsoft Excel 2015 and GraphPad Prism 8. Mean ± SEM is reported throughout the manuscript. Note that SEM for n=2 equals the range between the two points.

## Results

### Prdm12 exon V constitutive knockout mice show selective loss of the developing nociceptor population

To study the role of *Prdm12* during embryogenesis, we first generated homozygous null mice in which exon V was deleted from conception. This was done by crossing mice expressing an exon V floxed *Prdm12* allele (Fig. 1A-B) (Chen et al., 2020) to germline CAG-Cre mice (Sakai and Miyazaki, 1997), generating heterozygous *Prdm12*^*+/-*^ mice, which were then crossed to each other, producing *Prdm12*^*-/-*^ homozygotes. Using *in situ* hybridization with a probe specific for exon V, we confirmed that expression of this sequence was eliminated from mutant DRGs at E11.5 (Fig. 1C). *Prdm12*^*-/-*^ embryos appear grossly normal during development, and were observed to move and breathe normally immediately following cesarean section at E18.5, but newborn pups die within hours of birth. On closer examination, lumbar DRGs from *Prdm12*^*-/-*^ embryos were found to be smaller than those of control littermates (Fig. 1D). The relative size of mutant DRGs to control DRGs shrank from ∼68% at E11.5 to ∼43% at E18.5 (Fig. 1D).

**Figure 1.**
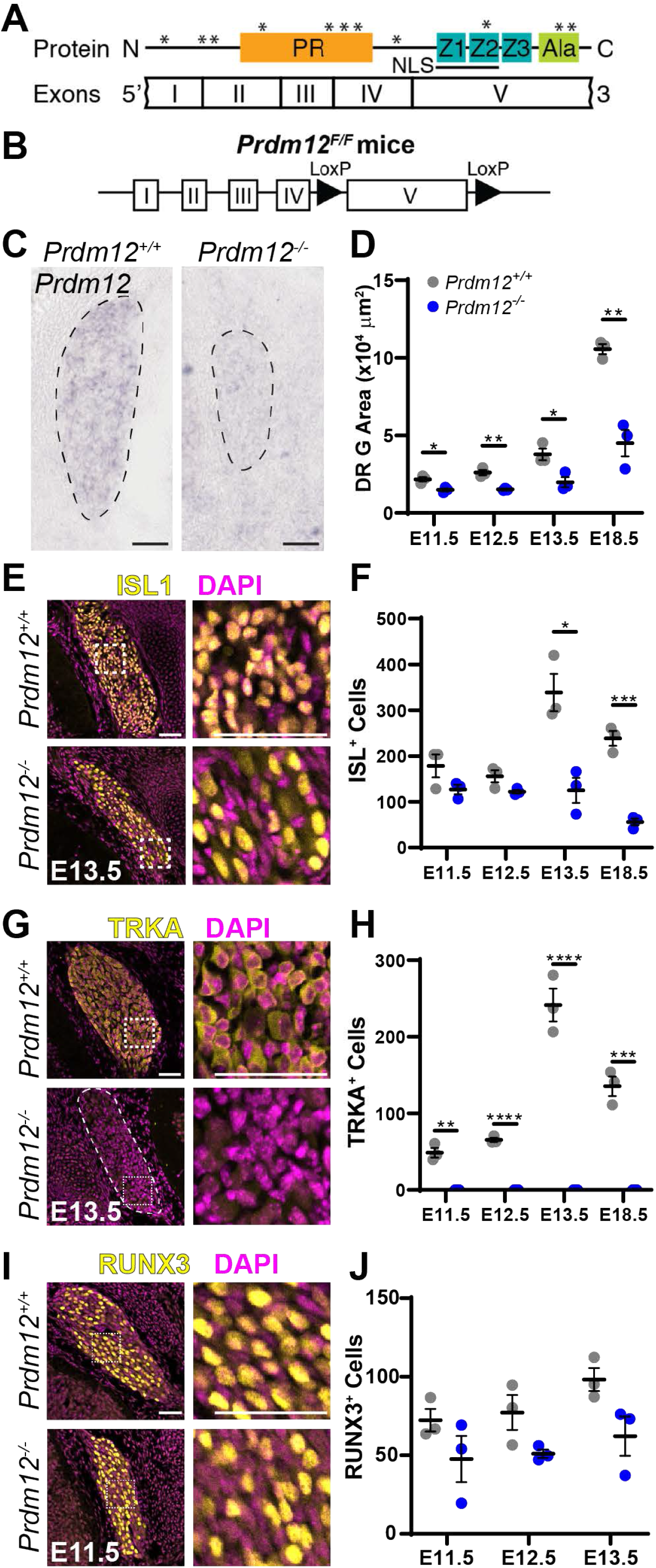
DRGs from *Prdm12*^*-/-*^ embryos are smaller and lack nociceptors. (A) PRDM12 protein domain structure with corresponding exons. *Human disease-causing mutations. (B) Schematic of the *Prdm12*^*F/F*^ allele. (C) *In situ* hybridization with an exon V-specific probe verified deletion of this transcript in *Prdm12*^*-/-*^ embryos. Scale bar 50 μm. (D) Quantification of DRG area from immunofluorescence images reveals *Prdm12*^*-/-*^ DRGs are smaller at all timepoints (E11.5 *p* = 0.017, E12.5 *p* = 0.003, E13.5 *p* = 0.023, E18.5 *p* = 0.003). (E) Representative images of ISL1 immunohistochemistry, with inset shown on right. Scale bars 50 μm. (F) Quantification reveals a similar number of ISL1^+^ cells at E11.5 and E12.5 in control and KO tissue, but a significant reduction in counts at E13.5 (*p* = 0.012) and E18.5 (*p* = 0.0005) in KO embryos. (G) Representative images of TRKA immunohistochemistry, with inset shown on right. Scale bars 50 µm. (H) Quantification reveals a complete absence of TRKA^+^ precursors in *Prdm12*^*-/-*^ embryos at all time points (E11.5 *p* = 0.001, E12.5 and E13.5 *p* < 0.0001, E18.5 *p* = 0.0004). (I) Representative images of RUNX3 immunohistochemistry, with inset shown on right. Scale bars 50 µm. (H) Quantification reveals no significant difference between control and KO DRGs at any timepoint. For all graphs, a data point represents the average across 3 DRGs taken from the lumbar region of a single embryo. Results are presented as mean ± SEM; statistical analysis performed with pairwise *t*-tests.

To identify what cellular changes occurred to result in smaller *Prdm12*^*-/-*^ DRGs, we analyzed how the number and types of neurons were affected. We performed immunohistochemistry with the pan-sensory neuron transcription factor, ISLET1 (ISL1), which defines differentiated sensory neurons, and TRKA, a specific marker for nociceptors (Fig. 1E-H) (Moqrich et al., 2004; Smeyne et al., 1994). We found that the number of neurons (ISL^+^) was unchanged at early embryonic time points (E11.5 and E12.5), but significantly decreased at later embryonic time points (E13.5 and E18.5) (Fig. 1E-F). In contrast, TRKA was completely absent at all timepoints in *Prdm12*^*-/-*^ mice, indicating the entire nociceptor lineage was lost. Notably, the number of ISL1^+^ and TRKA^+^ cells in control DRGs increases at E13.5, while remaining constant in KO embryos (Fig. 1F, H). Given that myelinated neurons are born before the unmyelinated neurons during DRG neurogenesis, we surmise that the sudden increase of ISL1^+^ and TRKA^+^ neurons in control tissue at E13.5 is due to the differentiation of the main pool of nociceptive neurons (Kitao et al., 2002; Lawson and Biscoe, 1979; Ma et al., 1999).

Because TRKA^+^ nociceptors were completely absent from the DRG at all embryonic time points, we wanted to test whether there was a compensatory increase in alternate cell fates. To investigate the effect of *Prdm12-*knockout on non-nociceptive sensory lineages, we stained DRGs for RUNX3 (runt-related transcription factor), which marks early proprioceptive neurons (Fig. 1I) (Inoue et al., 2002; Kramer et al., 2006; Levanon et al., 2002). While numbers of RUNX3^+^ cells trended lower in KO tissue at all timepoints, no significant differences between control and KO were noted. In fact, average RUNX3 levels were relatively constant from E11.5 to E13.5 (Fig. 1J) (Lallemend and Ernfors, 2012). Therefore, proprioceptor development is unaffected indicating that the fate of nociceptors do not switch to proprioceptors in *Prdm12*^-/-^ mice. Overall, our data suggest that absence of *Prdm12* during neurogenesis results in selective loss of the nociceptor population, while proprioceptive DRG neurons develop normally, consistent with reports in *Prdm12* exon II KO mice (Bartesaghi et al., 2019; Desiderio et al., 2019).

### Nociceptors fail to proliferate and differentiate in Prdm12^-/-^ embryos

We next sought to examine the developmental mechanism by which nociceptors are lost in *Prdm12*^*-/-*^ embryos. Two previous studies using a *Prdm12* exon II KO mouse suggest two possibilities: (1) nociceptors die by apoptosis (Desiderio et al., 2019), or (2) precursors fail to proliferate (Bartesaghi et al., 2019). To address these possibilities in our mouse model, we examined the changes in apoptosis using the marker cleaved caspase-3 and in proliferation using a thymidine analog (Fig. 2). We found that the total number of apoptosing cells was unchanged or even reduced at E12.5 in *Prdm12*^*-/-*^ embryos compared to controls (Fig. 2B). When normalized to DRG size, however, no significant differences were noted between control and *Prdm12*^*-/-*^ mice, suggesting that the overall rate of apoptosis was similar between groups (Fig. 2C). It is particularly notable that no difference was seen at E13.5, as an increase in apoptosis at this timepoint would specifically point to death of the newly ISL1^+^, TRKA^+^ cells normally present in controls (Fig. 1). Thus, it does not appear that nociceptors or their precursors are dying in increased numbers.

**Figure 2.**
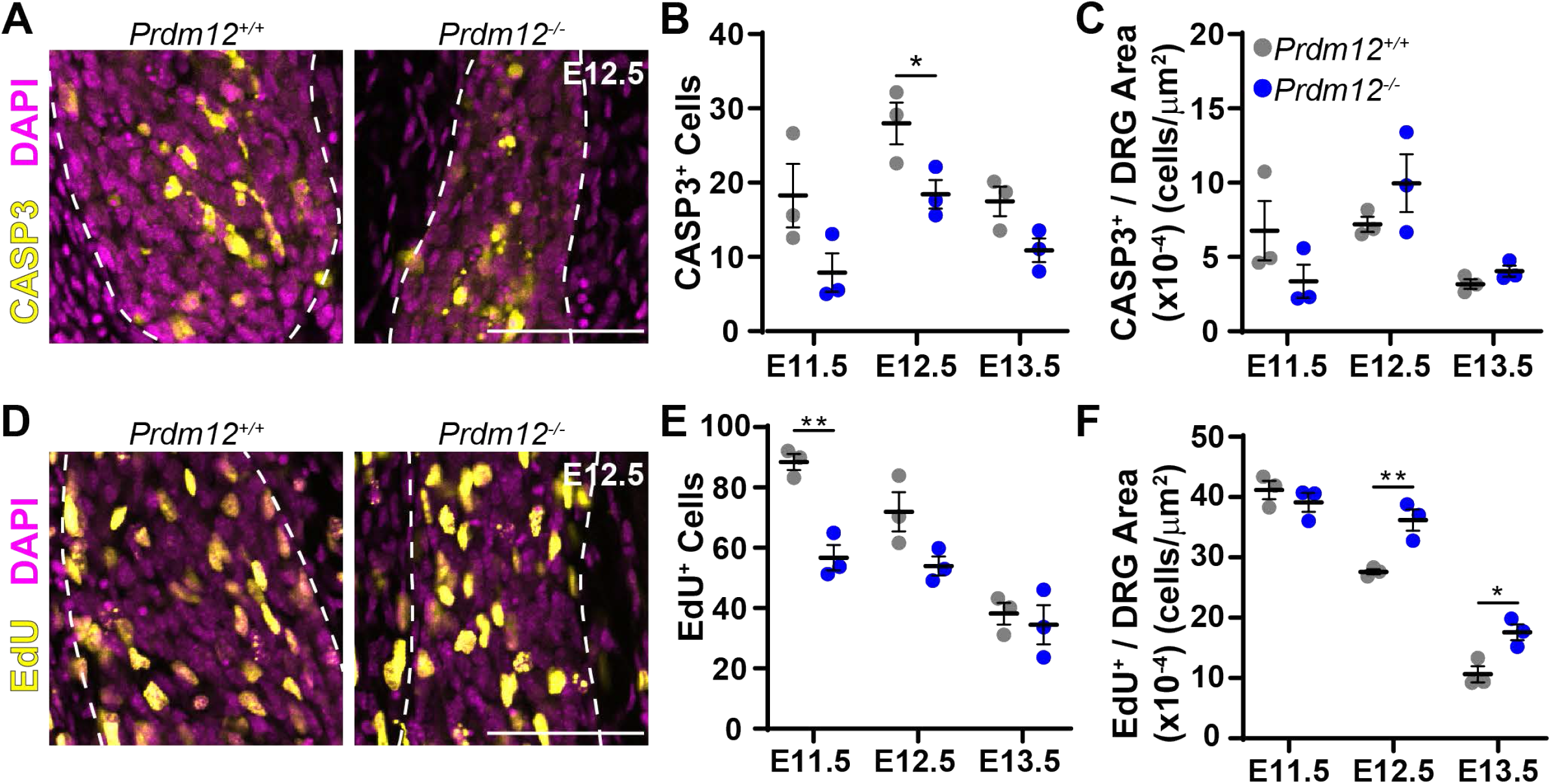
Defects in proliferation but not cell death in *Prdm12*^*-/-*^ mice. (A) Representative images of cleaved caspase-3 immunohistochemistry in E12.5 embryos. Scale bar 50 μm. (B) Quantification reveals a small but significant reduction of CASP3^+^ cells in *Prdm12*^*-/-*^ embryos at E12.5 (*p* = 0.043). (C) When normalized to DRG area, there is no significant difference in the number of CASP^+^ cells at any time point. (D) Representative images of DRGs labeled with EdU just prior to collection at E11.5. (E) Quantification reveals a significant reduction in EdU-labeled cells at E11.5 (*p* = 0.003) in *Prdm12*^*-/-*^ DRGs. (F) When corrected for DRG size, counts indicate increased relative EdU labeling in *Prdm12*^*-/-*^ DRGs at E12.5 (*p* = 0.009) and E13.5 (*p* = 0.021). For all graphs, a data point represents the average across 3 DRGs taken from the lumbar region of a single embryo. Results are presented as mean ± SEM; statistical analysis performed with pairwise *t*-tests.

We next looked at the effect of *Prdm12*-knockout on proliferation of sensory neuron precursor cells. To do this, we i.p. injected the thymidine analog 5-ethynyl-2’-deoxyuridine (EdU) into pregnant dams half an hour prior to collecting embryos to label proliferating cells at each time point. Using immunohistochemistry to visualize EdU-labeled cells, we found that the total number of EdU^+^ cells is significantly reduced in *Prdm12*^*-/-*^ DRGs at E11.5 (Fig. 2D-E). Thus, we infer that the progenitors that would make TRKA^+^ nociceptors are not present at E11.5, resulting in an overall decrease in the total number of proliferating cells (Fig. 2E). Furthermore, total EdU levels are similar in control and *Prdm12*^*-/-*^ DRGs at E12.5 and E13.5, indicating that non-nociceptive lineages are proliferating normally. When normalized to DRG size, there appears to be a significant increase in relative proliferation at E12.5 and E13.5 in *Prdm12*^*-/-*^ tissue, but this is because the DRGs are smaller due to the absence of nociceptors (Fig. 2F). Our findings are consistent with what Bartesaghi et al. describe in a constitutive *Prdm12* exon II knockout model showing a reduction in proliferation using phospho-histone H3 (pH3) staining (Bartesaghi et al., 2019). In summary, we have shown that *Prdm12* likely plays a role in the proliferation of progenitors that become nociceptors while proliferation of non-nociceptive populations is unchanged.

### Prdm12^AvilCKO^ mice have reduced cold and mechanical sensitivity, and pruriception

The above evidence, as well as the human phenotype of *Prdm12*-associated CIP suggest that loss of *Prdm12* results in insensitivity to pain due to a failure of development of nociceptive neurons (Chen et al., 2015). To test this hypothesis, we examined whether deletion of *Prdm12* leads to a painless phenotype. Because *Prdm12*^*-/-*^ mice die neonatally, we used a conditional knockout approach to specifically target sensory neurons using *Advillin*^*Cre/+*^ knockin mice (Hasegawa et al., 2007; Zhou et al., 2010). While *Advillin* protein is reported to be enriched in non-peptidergic isolectin B4 (IB4^+^) nociceptors (Hunter et al., 2018), our findings suggest the CRE recombinase in the *Advillin*^*Cre/+*^ knockin mice is broadly expressed in almost all DRG neurons, including those expressing *Prdm12* mRNA (see Fig. S1). Furthermore, although *Advillin* is reported to be expressed in other tissues, including endocrine cells, Merkel cells, and sympathetic ganglia (Hunter et al., 2018), *Prdm12* is not expressed in these tissues and thus, deletion in these tissues should not affect a nociceptive phenotype (Chen et al., 2015; Kinameri et al., 2008; Matsukawa et al., 2015; Nagy et al., 2015). We therefore crossed *Advillin*^*Cre/+*^ mice heterozygous for *Prdm12*^*-/+*^ to the *Prdm12*^*F/F*^ mice, and to a CRE-dependent fluorescent reporter (*R26*^*LSL-tdTomato*^, Ai14) (Madisen et al., 2010). The resulting *Prdm12*^*AvilCKO*^ mice survive, in contrast to the neonatal lethality seen with germline KO. *Prdm12*^*AvilCKO*^ were therefore tested to see whether loss of *Prdm12* specifically in DRGs causes deficits in pain sensation.

To investigate the phenotypic consequence of sensory neuron-specific *Prdm12*-knockout, we tested the sensation of *Prdm12*^*AvilCKO*^ mice to a variety of stimuli. Notably, mechanical nociception was reduced in mutant mice in both pinprick and tail clip assays (Fig. 3A-B). Nociceptive cold sensitivity to a 0-5°C cold plate was similarly reduced, reflecting that cold thermal-sensing nociceptors are affected as well (Fig. 3C). Curiously, while human *Prdm12*-associated CIP also causes heat insensitivity, thermal thresholds were unchanged using both the Hargreaves assay and hot plate at various temperatures (Fig. 3D-E), indicating that *Prdm12*^*AvilCKO*^ mice remain sensitive to nociceptive heat stimuli. As with cases of human CIP, non-nociceptive sensation remained intact (Chen et al., 2015). Responses of *Prdm12*^*AvilCKO*^ mice to low-threshold mechanical stimuli applied with von Frey hairs were similar to control, as were responses to dynamic stimuli applied with a brush stroke (Fig. 3F-G). Control and mutant mice also had a similar latency to response to a piece of tape applied to the plantar surface of the hindpaw, again reflecting normal tactile sensation (Fig. 3H). Finally, pruriception was tested by intradermal injection of either chloroquine or histamine into the nape of the neck. *Prdm12*^*AvilCKO*^ mice showed a significantly reduced scratch response to both stimuli, indicating that loss of *Prdm12* during development impacts itch sensation as well as nociception (Fig. 3I-J).

**Figure 3.**
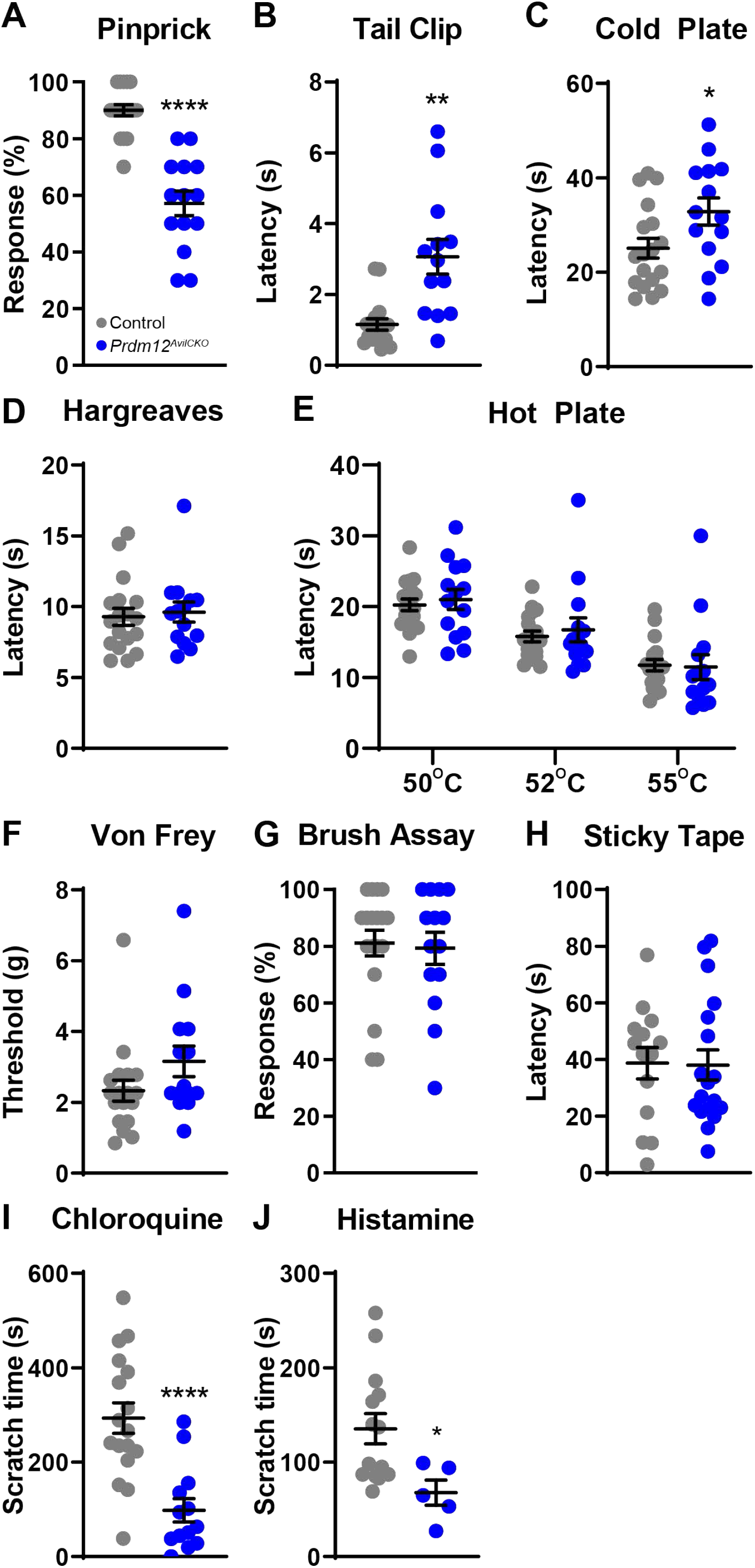
*Prdm12*^*AvilCKO*^ mice have reduced sensation to mechanical and cold nociceptive stimuli, and chemical pruritogens. (A-I) Panels show behavioral responses of *Prdm12*^*AvilCKO*^ mice (n = 14) and control littermates (n = 18); each group is split 50/50 M:F. (A) *Prdm12*^*AvilCKO*^ mice show reduced sensitivity to a sharp pin, *p* < 0.0001. (B, C) *Prdm12*^*AvilCKO*^ mice show a delayed response to a clip attached to the tail (B, *p* = 0.002) and when placed on a cold plate (C, 0-5°C, *p* = 0.033). (D, E) *Prdm12*^*AvilCKO*^ mice showed no differences in response to heat stimuli in the Hargreaves (D) or hot plate (E) assays. (F-H) Light touch is also normal in *Prdm12*^*AvilCKO*^ mice, which show similar withdrawal thresholds to von Frey hairs (F, *p* = 0.115), responses to dynamic light touch (G), and latency to removal of a piece of tape applied to the plantar surface of the hindpaw (H). (I) *Prdm12*^*AvilCKO*^ mice show reduced scratch time following intradermal chloroquine injection in the nape of the neck, *p* < 0.0001. (J) Scratch time is similarly reduced in *Prdm12*^*AvilCKO*^ mice (n = 5) compared to control littermates (n = 14) following intradermal histamine injection in the nape of the neck, *p* = 0.029. All behavioral data analyzed by *t*-test; results are presented as mean ± SEM.

### Developmental loss of Prdm12 reduces the number of nociceptors in the DRG

Having revealed a behavioral phenotype resulting from loss of *Prdm12*, we next assessed what molecular changes occurred in *Prdm12*^*AvilCKO*^ DRGs. To confirm that our genetic manipulation was successful, we performed RNAscope using a probe specifically targeting exon V of *Prdm12*. As expected, the probe detected mRNA transcript as multiple distinct puncta in control tissue, but not in mutant DRGs, indicating successful knockout of this region (Fig. 4I).

**Figure 4.**
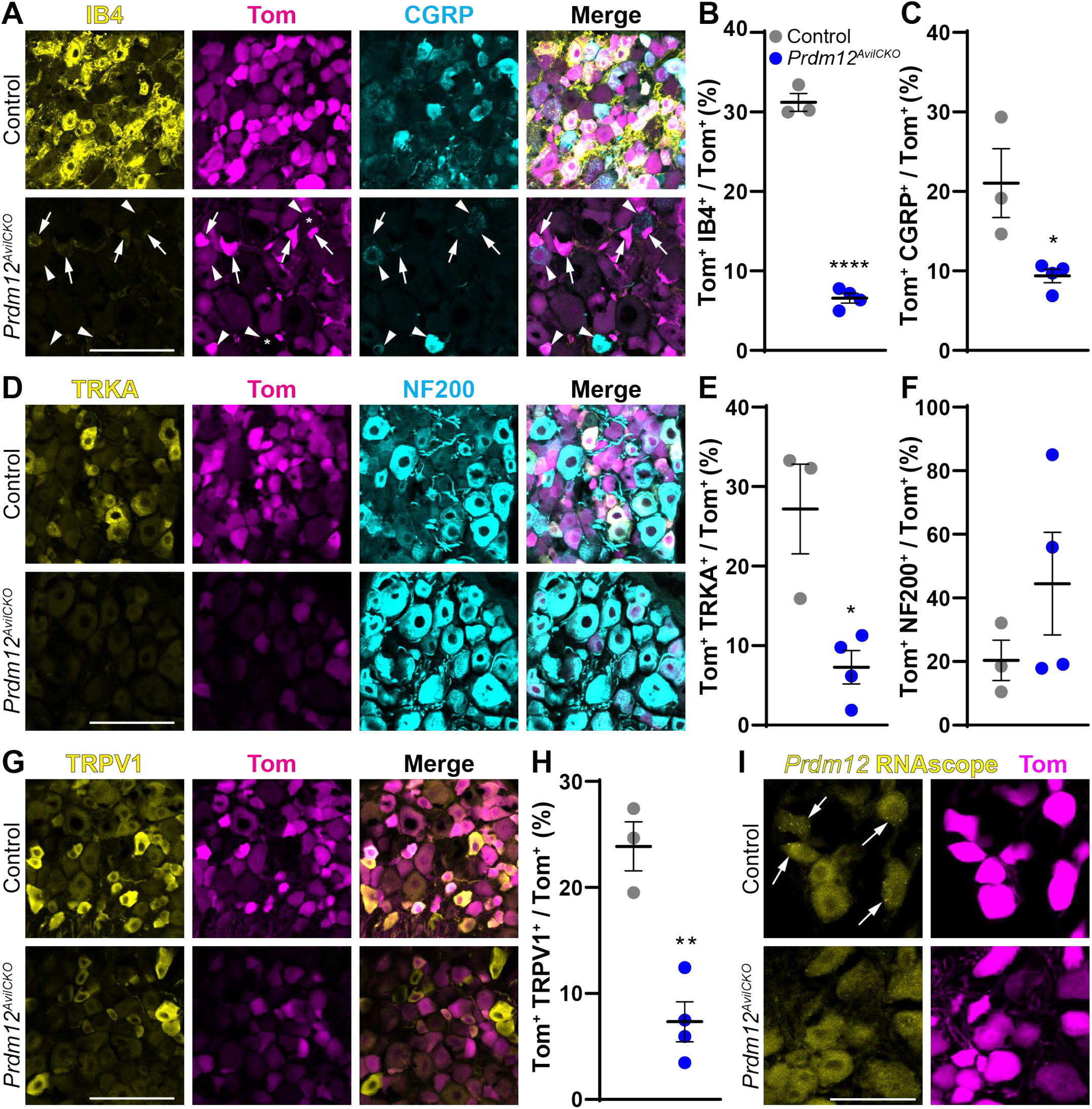
Nociceptor populations are reduced in *Prdm12*^*AvilCKO*^ mice. (A) Representative image showing number of IB4^+^ (arrows) and CGRP^+^ (arrowheads) nociceptors are reduced in *Prdm12*^*AvilCKO*^ DRGs. Scale bar 100 µm. (B, C) Quantification of IB4^+^ (B, *p* < 0.0001) and CGRP^+^ (C, *p* = 0.027) nociceptors as a percent of the total Tom^+^ population of sensory neurons. (D) Representative image of TRKA^+^ nociceptors and NF200^+^ myelinated neurons in *Prdm12*^*AvilCKO*^ and control mice. Scale bar 100 µm. (E) Quantification showing significant reduction of TRKA^+^ nociceptors, *p* = 0.0135. (F) Quantification of NF200^+^ neurons showing a wide range in *Prdm12*^*AvilCKO*^ mice, but no significant change from control littermates, *p* = 0.278. (G, H) Representative image (G) and quantification (H) of the reduction in number and intensity of TRPV1^+^ nociceptors, *p* = 0.0025. Scale bar 100 µm. (I) RNAscope using exon V-specific probes confirmed knockout of *Prdm12* from mutant DRGs. All analysis was completed using DRGs from lumbar levels 2 through 5; each data point represents the average count across 3 DRGs from control (n = 3) or *Prdm12*^*AvilCKO*^ (n = 4) mice taken after behavior analysis around 10 weeks of age. Scale bar 50 µm. All quantification analyzed by *t*-test; results are presented as mean ± SEM.

On further histological assessment, DRGs from *Prdm12*^*AvilCKO*^ mice were found to be missing the majority of their nociceptive neurons. The population of non-peptidergic IB4^+^ C-fibers was reduced by ∼80% in mutant DRG sections (Fig. 4A-B). The number of peptidergic CGRP^+^ neurons was similarly reduced by ∼50% (Fig. 4C). Unlike in the constitutive knockout (*Prdm12*^*-/-*^), *Prdm12*^*AvilCKO*^ mice still have TRKA^+^ neurons, though this population was reduced by ∼75% (Fig. 4D-E). With such a drastic loss of unmyelinated C-fibers, we expected the majority of cells remaining in the DRG to be myelinated. However, we found a wide variation in the percentage of NF200^+^ cells (18%-85%), with some mutant DRGs showing no change in the relative number of NF200^+^ cells (Fig. 4F). Finally, even though we saw no difference in heat sensitivity, TRPV1 was reduced by ∼70% (Fig. 4G-H). This raises the possibility that functional compensation in nociceptors expressing TRPM3 or TRPA1 remains unaffected by *Prdm12*-knockout, as these ion channels were shown to have an overlapping role in heat sensation with TRPV1 (Vandewauw et al., 2018). Furthermore, the retention of heat thermal nociception may be explained by timing of the knockout, which does not occur until E12.5 in mutant mice. Because *Prdm12* is expressed as early as E9.5, it is possible that the delay in knockout spares some populations of nociceptors (Chen et al., 2015; Kinameri et al., 2008; Sharma et al., 2020).

### Adult knockout of Prdm12 does not affect nociception in naïve or injured animals

While our results so far add to the field of knowledge regarding the role of *Prdm12* during nociceptor development, very little is known yet about the function of this transcription factor in adulthood. *Prdm12* continues to be expressed in nociceptors of mature DRGs (Fig. 4I) and through late adulthood (Chen et al., 2015; Kinameri et al., 2008; Sharma et al., 2020; Usoskin et al., 2014). We set out to investigate the adult role of *Prdm12* by crossing *Prdm12*^*F/F*^ mice with the *Avil*^*CreERT2*^*BAC* transgenic strain heterozygous for *Prdm12*^*-/+*^ (Lau et al., 2011) to generate *Prdm12*^*AvilERT2CKO*^ mice. The *Cre*^*ERT2*^ allows for temporal control of recombination, as it only becomes nuclear localized with exposure to tamoxifen.

To test whether loss of *Prdm12* from mature nociceptors affects pain sensation, *Prdm12*^*AvilERT2CKO*^ mice and control littermates were injected with tamoxifen at 8 weeks of age to delete *Prdm12* from sensory neurons (Fig. 5A). After four weeks, baseline nociceptive responses to von Frey, cold plantar assay (CPA), and Hargreaves were used to assess mechanical, cold, and heat nociception, respectively. No differences in responses were found between control and mutant mice with any of these assays (Fig. 5, “BL”). Similarly, no difference was found in the licking response to capsaicin injected into the hindpaw, suggesting TRPV1^+^ nociceptors were functioning normally (Fig. 5B).

**Figure 5.**
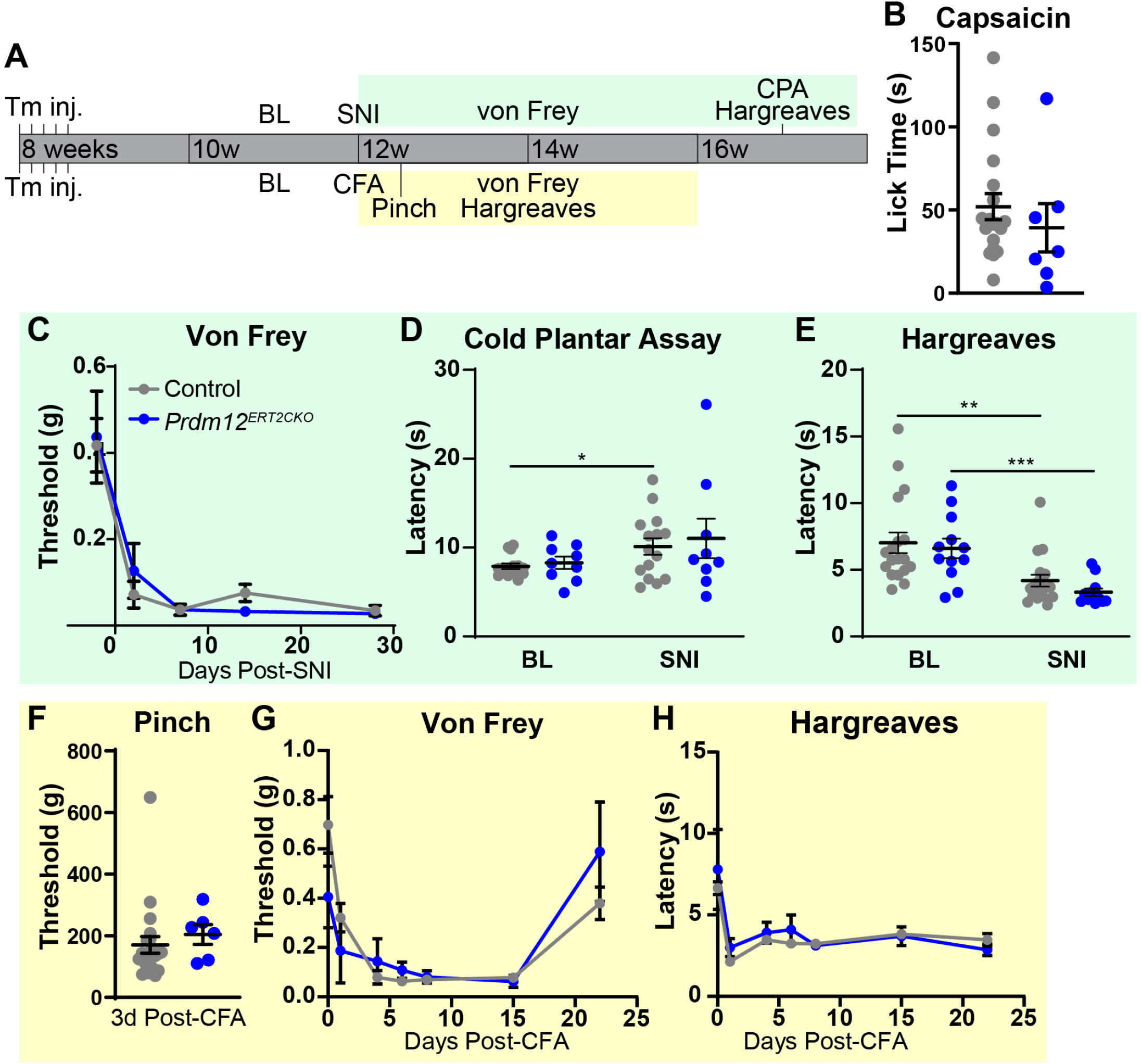
Knockout of *Prdm12* in adulthood does not reduce pain sensitivity in naïve or injured mice. (A) Schematic showing experimental timeline. (B) No difference was observed in the time spent licking after capsaicin injection into the hindpaw between *Prdm12*^*AvilERT2CKO*^ (n = 7, 3:4 M:F) and control (N=19, 11:8 M:F) mice. (C-E) Behavioral results before and after SNI. (C) Time course of withdrawal thresholds for *Prdm12*^*AvilERT2CKO*^ (n = 12, 5:7 M:F) and control (n = 18, 8:10 M:F) mice showing both groups developed mechanical allodynia following SNI. (D) Responses of *Prdm12*^*AvilERT2CKO*^ mice (n = 9, 5:4 M:F) to cold plantar assay did not differ significantly from control (n = 15, 7:8 M:F) at baseline or 4 weeks post-SNI. Control mice did show a slight increase in latency to response following SNI, *p* = 0.038. (E) Both groups experienced heat hyperalgesia four weeks post-SNI, but did not differ from each other at either time point. Same n as (C), control *p* = 0.004, *Prdm12*^*AvilERT2CKO*^ *p* = 0.0009. (F-H) Behavioral results following CFA injection. (F) No difference was observed in the withdrawal threshold to paw pinch between *Prdm12*^*AvilERT2CKO*^ (n = 6, 3:3 M:F) and control (n = 21, 12:9 M:F), which was tested 3 days after CFA injection. Note that withdrawal thresholds are several-fold higher due to the larger area over which pressure is applied with the rodent pincher compared to von Frey filaments. (G) Time course of withdrawal thresholds showing both groups developed tactile allodynia following CFA, and recovered over the same time period. Same n as (F). (H) Both groups developed heat hyperalgesia following CFA injection. Same n as (F). Statistical analysis by 2-way ANOVA for (C), (G), (H); pairwise t-tests for other data sets.

Although we found no changes in nociception in *Prdm12*^*AvilERT2CKO*^ mice, PRDM12 has been implicated in chromatin remodeling complexes with EHMT2 (Yang and Shinkai, 2013). EHMT2 has been found to mediate mechanical allodynia and heat hyperalgesia upon nerve injury through repression of voltage-gated potassium channels (Laumet et al., 2015; Liang et al., 2016). Therefore, we hypothesized that PRDM12 may also play a role in sensitization following injury through its interactions with EHMT2. We assessed the effect on allodynia and hyperalgesia in *Prdm12*^*AvilERT2CKO*^ mice in the setting of both nerve and inflammatory injury.

Spared nerve injury was performed on *Prdm12*^*AvilERT2CKO*^ mice and littermate controls 4 weeks after tamoxifen injection, and nociceptive behavior was reassessed. Von Frey testing was performed at several timepoints to establish a time course of responses. Surprisingly, no reduction in mechanical allodynia was found, as both control and *Prdm12*^*AvilERT2CKO*^ mice became hypersensitive following SNI, with no difference in paw withdrawal thresholds between the groups at any time point (Fig. 5C). Neither controls nor mutants developed cold allodynia following SNI, as measured with the cold plantar assay. This is likely because the posture adopted by injured mice lifts the hypersensitized region away from the glass, preventing direct application of the cold stimulus. In fact, controls showed a slight increase in latency to response (Fig. 5D). Both groups of mice experienced similar degrees of heat hyperalgesia measured by Hargreaves, but again no difference emerged between controls and mutants (Fig. 5E). These findings indicate that *Prdm12* is does not affect hypersensitivity of mature nociceptors following nerve injury.

We next wanted to determine whether loss of *Prdm12* would confer protection following inflammatory injury. To test this, we injected CFA into the hindpaw of control and mutant mice, and measured the responses to mechanical and thermal stimuli over the next three weeks (Fig. 5F-H). Again, no differences in responses were noted between controls and mutants. Both groups developed similar levels of tactile allodynia, which faded approximately three weeks after injection (Fig. 5F-G). As an additional measure, we also tested sensitivity to pinch in the inflamed paw 3 days after CFA injection, and found no difference between cohorts (Fig. 5F). Similarly, no differences in thermal hyperalgesia were noted between the two groups (Fig. 5H). Overall, our results show that loss of *Prdm12* function during adulthood does not alter pain sensation or affect the development of allodynia and hyperalgesia following injury.

### Molecular and transcriptional changes following Prdm12-knockout in adult

To look for molecular changes that may illuminate the observed lack of phenotype, we next used immunohistochemistry to assess for any changes in the DRG. Lumbar DRGs were taken from control and *Prdm12*^*AvilERT2CKO*^ mice 4 weeks post-SNI, so that we could investigate changes following both tamoxifen injection and neuropathic injury. We first verified successful knockout of *Prdm12* using the exon V-specific RNAscope probe, and found it be absent from *Prdm12*^*AvilERT2CKO*^ DRG neurons (Fig. 6A). Assessment of nociceptor populations within the DRG, however, found no significant differences in levels of IB4, CGRP, TRKA, NF200, or TRPV1 between groups (Fig. 6B-G). Additionally, no significant changes in these markers were noted between DRGs ipsilateral and contralateral to SNI, suggesting that nerve injury also does not alter the numbers of these nociceptor populations (Fig. 6B-G), consistent with the subtle transcriptional changes of these markers upon injury (Renthal et al., 2020).

**Figure 6.**
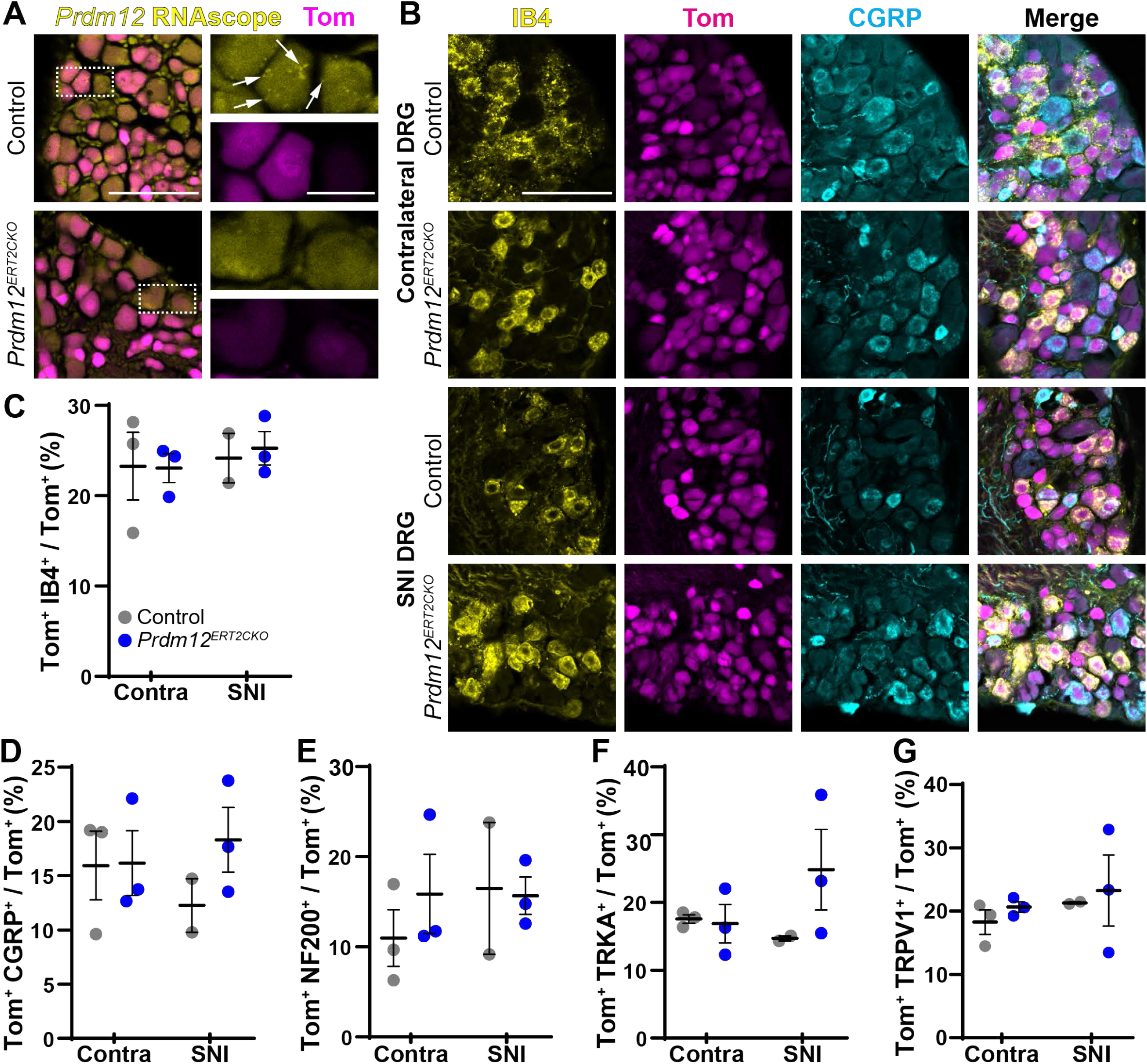
DRG nociceptor populations are unchanged following *Prdm12* knockout and/or SNI. (A) Exon V-specific RNAscope verified loss of mRNA transcript in *Prdm12*^*AvilERT2CKO*^ mice. Scale bar 100 μm. Inset arrows indicate mRNA puncta detected by the probe; inset scale bar 25 μm. (B) Representative images of lumbar DRGs from control and *Prdm12*^*AvilERT2CKO*^ DRGs contralateral to and ipsilateral to SNI with immunohistochemistry for IB4 and CGRP. Scale bar 100 μm. (C-G) Quantification of these images revealed no changes in the number of IB4^+^ (C), CGRP^+^ (D), NF200^+^ (E), TRKA^+^ (F), or TRPV1^+^ (G) neurons after SNI in either control or *Prdm12*^*AvilERT2CKO*^ mice, or between these two groups at either time point. Each data point represents the average count across 3 DRGs taken from the L2-L5 region of n = 2 control mice post-SNI, or n = 3 mice for all other conditions. DRGs were collected after behavior assessment, at 18 weeks. Graphs show mean ± SEM; statistical analysis with pairwise t-tests.

To investigate broader transcriptional changes that occur following loss of *Prdm12*, we performed bulk mRNA-seq of DRGs harvested from *Prdm12*^*AvilERT2CKO*^ and control mice two weeks after tamoxifen injection. SNI was not performed on these animals. As expected, exon V is specifically knocked out in *Prdm12*^*AvilERT2CKO*^ mice (Fig. 7A). Surprisingly, though, we found that the majority of the differentially expressed genes (DEGs) were decreased in knockout mice, suggesting PRDM12 acts as a transcriptional activator (Fig. 7B). This is in contrast to prior evidence suggesting that histone methyltransferase activity associated with *Prdm12* is repressive (Kinameri et al., 2008; Thelie et al., 2015).

**Figure 7.**
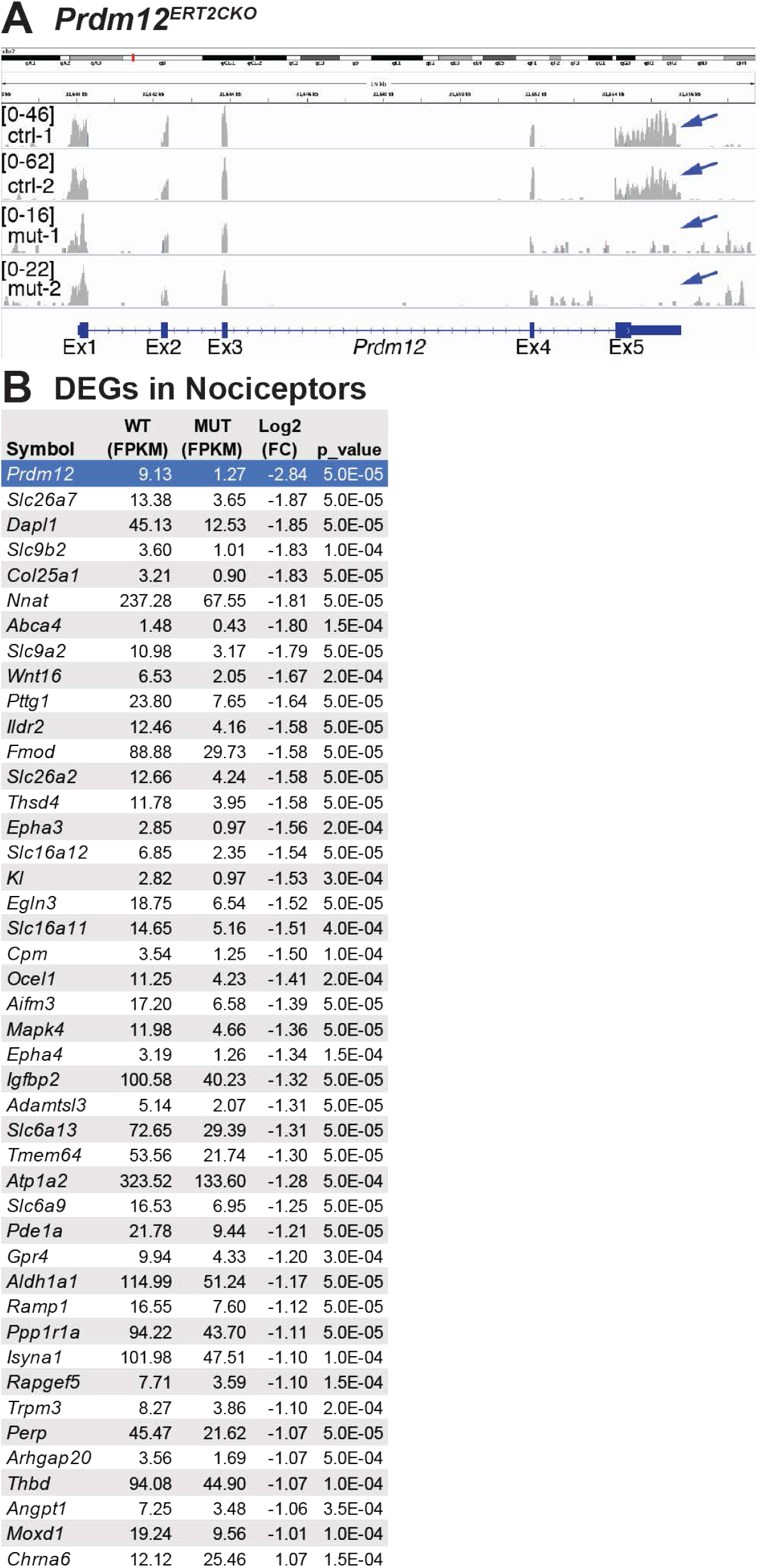
Transcriptional changes in *Prdm12*^*AvilERT2CKO*^ mice. (A) Sequencing reads from two control (ctrl-1 and 2) and two mutant (mut-1 and 2) samples show that exon V is specifically knocked out (arrows). (B) 43 genes expressed in nociceptors are decreased in the mutant (negative log2 (Fold Change)), and 1 gene is increased (*Chrna6*). FPKM values are averaged across both samples.

Because mRNA was harvested from all cells in the DRG, we cross-referenced the initial list of 150 DEGs to a published data set from scRNA-seq of DRG neurons (Sharma et al., 2020) (Fig. 7B, Supplemental Table 1). 44 of these genes were nociceptor-specific—43 were decreased after loss of *Prdm12*, and only 1 gene, *Chrna6*, was increased (Fig. 7B). Most of the nociceptive genes identified in our data set are not classically used to define nociceptor cell types. One exception is *Trpm3*, which has been shown to mediate nociceptive heat sensation. Notably, two other ion channels involved in nociceptive heat sensation, *Trpv1* and *Trpa1*, were not changed in our dataset (Vandewauw et al., 2018). Overall, despite the lack of overt differences in pain sensation in *Prdm12*^*AvilERT2CKO*^ mice, it appears that loss of function of *Prdm12* does have a role in transcriptional control of adult nociceptors, but the exact nature of that role requires further investigation.

## Discussion

In this study, we explore the role of *Prdm12* both during embryonic development and in the adult. We find that while it is necessary for nociceptor neurogenesis, its function in mature nociceptors is unclear. We find that constitutive deletion of *Prdm12* exon V precludes development of the entire nociceptor lineage, normally marked by TRKA expression, resulting in smaller DRGs with fewer differentiated neurons. Moreover, our findings suggest that this is in part due to reduced precursor proliferation. Furthermore, we show that in a conditional knockout model, mice have reduced sensitivity to certain modalities of pain and itch, with correspondingly reduced nociceptor populations in the DRGs, but curiously heat sensation is spared. Lastly, we provide evidence that the function of *Prdm12* differs in adult DRGs compared to neurogenesis, as knockout did not significantly alter pain phenotype or nociceptor populations even in the context of injury and inflammation.

### Prdm12 is required for nociceptor development

We find that constitutive deletion of *Prdm12* exon V is sufficient to impede nociceptor development. As with prior reports using an exon II-knockout, we find that expression of TRKA is completely absent from DRGs of *Prdm12*^*-/-*^ embryos at all embryonic time points, supporting a key role in specification of this lineage (Bartesaghi et al., 2019; Desiderio et al., 2019). Our findings indicate that nociceptors fail to develop in *Prdm12*^*-/-*^ mice likely due to defects in specification and/or proliferation of progenitors that are likely to become nociceptors, and not due to an increase in cell death or respecification to proprioceptors, consistent with findings by Bartesaghi et al.

### Conditional knockout of Prdm12 from sensory neurons recapitulates aspects of human CIP

Here we show that *Prdm12*^*AvilCKO*^ mice bred to selectively remove *Prdm12* from DRGs during neurogenesis have deficiencies in several sensory modalities. Specifically, they have reduced mechano-nociception and cold nociception, as well as pruriception, all of which are hallmarks of CIP (Indo, 2014). These behavioral differences were accompanied by losses of IB4^+^, CGRP^+^, TRKA^+^, and TRPV1^+^ nociceptors from the DRGs. These changes bear a striking resemblance to the phenotype observed following ablation of Nav1.8^+^ postmitotic sensory neurons (Abrahamsen et al., 2008). These mice also show reduced IB4^+^ and CGRP^+^ nociceptor populations, as well as reduced TRKA and TRPV1 expression, and defects in cold and mechanical nociception, but not heat. These similarities raise the possibility that knockout of *Prdm12* in our model had a greater impact on Nav1.8^+^ neurons than the nociceptor population as a whole, leading to residual pain sensation. Interestingly, though, both Nav1.8-expressing nociceptor-ablated mice and *Trpv1*^*-/-*^ mice, which also have no heat phenotype at baseline, show reduced heat hyperalgesia following inflammatory injury (Abrahamsen et al., 2008; Davis et al., 2000), a phenotype not assessed in the *Prdm12*^*AvilCKO*^ mice.

### Residual pain in Prdm12^AvilCKO^ and Prdm12^AvilERT2CKO^ mice may indicate autonomous PR domain function

Comparing and contrasting the phenotype between the *Prdm12* exon II (Bartesaghi et al., 2019; Desiderio et al., 2019) and exon V (shown here) knockout mouse models can give us some clues as to the function of different domains within the PRDM12 protein. In the exon II knockout, exons II-V, which code for all functional domains for PRDM12 including the PR domain, are deleted (Bartesaghi et al., 2019; Desiderio et al., 2019). In the exon V knockout, the coding sequence for the three zinc finger domains, polyalanine repeat, and a nuclear localization sequence, are deleted. We found in our adult *Prdm12*^*AvilERT2CKO*^ mice that transcripts for exons I-IV, which includes the PR domain, are still expressed albeit at reduced levels (∼35%) (Fig. 7A). Therefore, while the putative interaction with EHMT2 that occurs through ZnF2 is disrupted in both the exon II and exon V knockout mouse models (Yang and Shinkai, 2013), the PR domain is potentially still expressed in the *Prdm12* exon V mouse model.

At the phenotypic level, subtle differences are seen in the *Prdm12* exon II and exon V sensory neuron-specific conditional knockout mice. In the knockout of *Prdm12* exon II, also using the *Avil*^*Cre/+*^ strain, mice were observed to develop eye opacities, as well as tail and facial scratches and wounds, similar to clouding of the eye and self-mutilating injuries seen in human CIP patients (Chen et al., 2015; Desiderio et al., 2019), but no such phenotype was noted in the *Prdm12*^*AvilCKO*^ mice presented here. It is possible that the PR domain is translated in the *Prdm12*^*AvilCKO*^ mice, carrying out an as-yet undescribed function independent of EHMT2, leading to a less severe phenotype than that seen with the sensory-specific conditional knockout of *Prdm12* exon II (Desiderio et al., 2019). Strain differences could also be another variable since our model was on a mixed background, while the exon II KO was in C57Bl/6J mice. Furthermore, the retention of exons I-IV in the adult *Prdm12*^*AvilERT2CKO*^ mice might contribute to the lack of detectable nociceptive phenotype in the adult. Further studies directly comparing the nociceptive behaviors in the exon II and exon V conditional KO mice will lend further insight into the potential role of the PR domain.

### Prdm12 may function as both an activator and a repressor, with distinct functions in adult nociceptors

As a whole, the PRDM family of transcription factors can have multifaceted roles including being an activator of cellular lineages and repressor of alternative fates, even within the same cell (Hohenauer and Moore, 2012). Early evidence suggested PRDM12 functions primarily as a repressor, due to its interactions with EHMT2, and its role in repressing neighboring progenitor domains of the V1 progenitor population in the developing spinal cord (Fog et al., 2012; Kinameri et al., 2008; Thelie et al., 2015). However, the RNA-sequencing data we report here points to PRDM12 serving as a transcriptional activator in the adult context, given that almost every transcript identified was downregulated in *Prdm12*^*AvilERT2CKO*^ mice. While it is possible that PRDM12 could be repressing a repressor, we are unable to identify a repressor that is upregulated following *Prdm12*-knockout. Furthermore, overexpression of PRDM12 with NEUROG1 in *Xenopus laevis* explants induces expression of several other genes essential for sensory neurogenesis, such as TRKA, while germline knockout in mice resulted in both increases and decreases of downstream targets during development (Desiderio et al., 2019; Nagy et al., 2015). Evidence thus suggests that PRDM12 may act as either a repressor or activator, and that this activity may change over developmental time, with PRDM12 being an activator in the adult mouse.

It is also notable that with the exception of *Prdm12* itself, there was almost no overlap in DEGs identified in our work and by Desiderio et al. Developmentally, *Prdm12* regulates an array of genes involved in generation of spinal cord interneurons, as well as transcriptional regulators of sensory neuron differentiation (Desiderio et al., 2019). DEGs identified in our data set are not obviously related to nociceptor neurogenesis, nor are they generally used to define nociceptor cell types, however some are nociceptor-specific (Sharma et al., 2020). Given that PRDM12 is proposed to induce *Ntrk1* expression through interactions with NEUROG1, which is only present during DRG embryogenesis, the lack of overlap between datasets is not surprising, but again points to alternate roles of *Prdm12* during development and in adult DRGs.

Indeed, the role of *Prdm12* in adult DRGs remains elusive. We found that exon V deletion in 8-week-old *Prdm12*^*AvilERT2CKO*^ mice did not affect a nociceptive phenotype under baseline, inflammatory, or neuropathic conditions. This was surprising for two reasons. Firstly, as already described, PRDM12 is thought to interact with EHMT2. The latter methyltransferase is normally upregulated after nerve injury, and developmental knockout or inhibition of EHMT2 reduces tactile allodynia and thermal hypersensitivity after neuropathic injury (Laumet et al., 2015; Liang et al., 2016). We hypothesized that PRDM12 would play a role in sensitization following nerve injury through its interactions with EHMT2, but this was refuted by the lack of a difference in sensitivity in *Prdm12*^*AvilERT2CKO*^ mice following SNI. As EHMT2 and PRDM12 proteins interact via ZnF2, which is deleted in our model, our data suggests that PRDM12 does not play a role in EHMT2-dependent hypersensitivity. Second, overexpression of *Chrna6*, which encodes the α6 subunit of the nicotinic acetylcholine receptor (nAChR), is protective against tactile allodynia in both neuropathic and inflammatory injury models (Wieskopf et al., 2015). *Chrna6* was also the only gene found to be upregulated in *Prdm12*^*AvilERT2CKO*^ mice, about two-fold over control levels. Given the absence of a phenotype in our mice, it is possible that a further increase in expression level is required to achieve protection against allodynia.

It is clear that *Prdm12* plays an integral role in the development of sensory neurons, and further study is needed to clarify the precise mechanisms requiring PRDM12 that specify the nociceptive lineage. Beyond development, however, *Prdm12* remains a specific marker of nociceptive neurons. While our study demonstrates that direct inhibition of its activity may not provide analgesic relief, we do find transcriptional changes resulting from its loss, and contend that further study is needed to define the precise nature of these changes, and whether they present a different angle for development of novel analgesics.

## Acknowledgements

This work was supported by F31 NS111796 and the William F. and Grace H. Kirkpatrick Award to M.A.L., NIH R01 DK114036 to C.L., Rita Allen Foundation Award in Pain, Welch Foundation, President’s Research Council Award, and Kent Waldrep Foundation to H.C.L. We thank Thomas Jessell for the RUNX3 antibody, Tou Yia Vue for help with EdU staining, Fan Wang for the *Advillin*^*Cre/+*^ mice, the UTSW Microarray Core Facility, UTSW Transgenic Core Facility, Stephanie Shiers, Rahul K. Kollipara, and the Johnson lab for technical assistance, Saida Hadjab for helpful discussions, Ted Price and Jane Johnson for critical reading of the manuscript.

## Author Contributions

H.C.L. designed and supervised the study. M.A.L. performed most experiments; M.G. and K.M.C. assisted with immunostaining experiments and microscopy analysis. C.L. generated and provided the *Prdm12*^*F/F*^ mice prior to publication. M.A.L. and M.G. prepared the figures. M.A.L. and H.C.L. wrote the paper with input from all other authors.

---

**Supplemental Figure 1.**
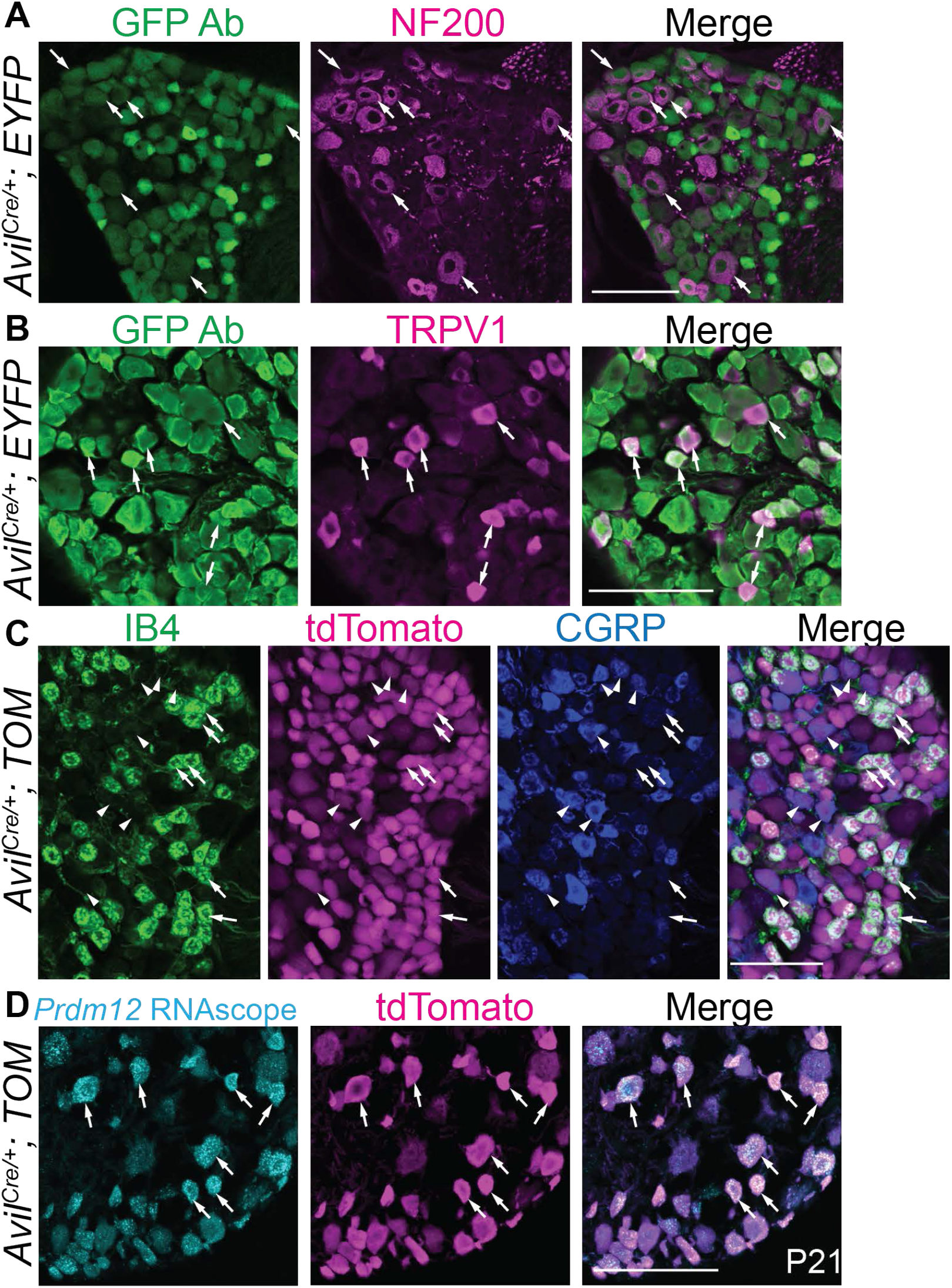
Characterization of a sensory neuron specific CRE line. (See Fig. 3). *Avil*^*Cre/+*^ crossed to a CRE-dependent reporter identifies DRG neurons expressing CRE recombinase. (A-C) *Avil*^*Cre/*+^-lineage neurons colocalize with myelinated DRG neurons (A, NF200^+^, arrows) unmyelinated TRPV1^+^ neurons (B, arrows), and nonpeptidergic (C, IB4^+^, arrows) and peptidergic (C, CGRP^+^, arrowheads) C-fibers. (D) The *Avil*^*Cre/*+^-lineage also colocalizes with *Prdm12* mRNA expression (arrows, RNAscope). Scale bars 100 μm.

